# Dynamic regulation of the *Trypanosoma brucei* transferrin receptor in response to iron starvation is mediated *via* the 3’UTR

**DOI:** 10.1101/441931

**Authors:** Corinna Benz, Winston Lo, Nadin Fathallah, Ashley Connor-Guscott, Henry J. Benns, Michael D Urbaniak

## Abstract

The bloodstream form of the parasite *Trypanosoma brucei* obtains iron from its mammalian host by receptor-mediated endocytosis of host transferrin through its own unique transferrin receptor (*Tb*TfR). Expression of *Tb*TfR rapidly increases upon iron starvation by post-transcriptional regulation through a currently undefined mechanism that is distinct from the mammalian iron response system. We have created reporter cell lines by fusing the *Tb*TfR 3′UTR or a control Aldolase 3’UTR to reporter genes encoding GFP or firefly Luciferase, and inserted the fusions into a bloodstream form cell line at a tagged ribosomal RNA locus. Fusion of the *Tb*TfR 3’UTR is sufficient to significantly repress the expression of the reporter proteins under normal growth conditions. Under iron starvation conditions we observed upregulation of the *Tb*TfR 3’UTR fusions only, with a magnitude and timing consistent with that reported for upregulation of the *Tb*TfR. We conclude that the dynamic regulation of the *T. brucei* transferrin receptor in response to iron starvation is mediated *via* its 3’UTR, and that the effect is independent of genomic location.

## Introduction

The obligate extracellular parasite *Trypanosoma brucei* has a complex digenetic lifecycle between a tsetse fly vector and a range of mammalian hosts. The bloodstream form of *T. brucei* has evolved a unique transferrin receptor (*Tb*TfR) that allows it to obtain the essential element iron through receptor mediated endocytosis of host transferrin (Tf) (1, 2). Different *Tb*TfR genes encode proteins with varying affinities for Tf from different mammals, and the occurrence of multiple *Tb*TfR genes has been suggested to allow the parasite to adapt to a wide host range (3, 4).

A subset of *T. brucei* genes are transcribed by RNA polymerase I (RNA pol I), including the essential Variant Surface Glycoprotein (VSG) which forms a dense surface coat that enables the parasite to evade the host’s innate and adaptive immune responses, and which undergoes antigenic variation from a repertoire of ∼1500 *VSG* genes (5). Of the 15 subtelomeric VSG bloodstream expression sites (BES), only one is active at a time so that a single *VSG* is transcribed from a discrete location within the nucleus (6). Antigenic variation requires replacement of the *VSG* within the active BES or a switch to a different BES, with the latter also causing a change in the expressed complement of expression site associated genes (*ESAGs*) that are located on the *VSG* polycistronic transcriptional unit. The *VSG* promoter proximal genes *ESAG6* & *ESAG7* form the heterodimeric *Tb*TfR (1), which is evolutionarily distinct from mammalian TfR and structurally resembles a truncated VSG homodimer (7). Both monomers of *Tb*TfR are extensively *N*-glycosylated and are membrane associated via the GPI anchor present on ESAG6 (8).

Under basal conditions only 3 × 10^3^ *Tb*TfR heterodimers (1) are expressed despite *ESAG6 & 7* being located on the same polycistronic transcriptional unit as the highly abundant VSG (5 × 10^6^ homodimers). Under iron starvation conditions expression of the *Tb*TfR rapidly increases equally at the mRNA and protein level, with lack of increase in *VSG* mRNA suggesting that regulation occurs through a currently undefined post-transcriptional mechanism (9, 10). Reducing the uptake of iron using the iron chelator deferoxamine, culturing with different mammalian serum, incubation with anti-TfR antibodies, or competition with apo-Tf all result in a rapid 2.5 – 5-fold upregulation of the *Tb*TfR and a corresponding increase in Tf uptake (4, 9–11). Interestingly, *Tb*TfR upregulation occurs before intracellular iron stores are depleted and cells continue to divide for 48 h, suggesting that cells are responding to changes in iron flux. The mechanism is distinct from the post-transcriptional Iron Response Element (IRE) / Iron Response Protein (IRP) system found in mammals, as knockout of the *T. brucei* IRP-1 homologue aconitase has no effect on *Tb*TfR regulation (9). As *Tb*TfR is a multi-gene family that occurs in an atypically regulated locus, and available antibodies cross-react with different ESAG6 & 7 glycoproteins, the direct study of *Tb*TfR regulation is challenging.

The BES commonly active in *T. brucei* bloodstream form culture adapted strains (BES1, expressing VSG221) contains *ESAG6 & 7* genes with nanomolar affinity for bovine Tf, but that binds canine Tf only poorly (3, 4). Prolonged iron starvation (>7 days) induced by changing from growth in media supplemented with bovine serum to canine serum selects for cells that have altered the identity of the expressed *ESAG6 & 7* and *VSG*, either by switching to another BES or replacing the genes in the active BES1 (4, 11). Switching events can be prevented by supplementing the canine serum with bovine Tf, demonstrating that the adaption is driven by iron starvation (3). Under normal physiological conditions the concentration of available host Tf is unlikely to limit trypanosome growth, but uptake may become limiting in later stages of an infection when competition with anti-*Tb*TfR antibodies and/or host anaemia come into consideration (11, 12).

There is mounting evidence of the importance of the 3’UTR in the post-transcriptional regulation of developmentally regulated genes in *T. brucei*. The stage-specific regulation of both RNA pol I transcribed VSG (13) and procyclin (14), and the RNA pol II transcribed COX genes (15) has been demonstrated to occur, at least in part, due to recognition of motifs within their respective 3’UTRs. The developmental regulation of *ESAG9* depends on a 34-nucleotide bifunctional element in the 3’UTR that confers both positive and negative regulation (16), and an RNA binding protein that negatively regulates *ESAG9* has recently been identified through a genome-wide RNA interference screen (17). Here, we investigate the importance of the *Tb*TfR 3’UTR in the dynamic regulation of *Tb*TfR in response to iron starvation using a simplified reporter system. By fusing the *ESAG6*-3’UTR to reporter genes encoding GFP or firefly Luciferase (fLUC), we demonstrate that the 3’UTR alone is sufficient to confer dynamic regulation of gene expression in response to iron starvation.

## Material and methods

### Cell lines

The culture adapted monomorphic *T. brucei brucei* Lister 427 bloodstream form 2T1 cell line (18), containing a tagged RRNA locus, were cultured in HMI-11T (19) containing 0.2 μg/mL Puromycin and 0.5 μg/mL Phleomycin at 37 °C in a 5% CO_2_ incubator. Transfected 2T1 cell lines were selected and maintained with 2.5 μg/mL Hygromycin in place of Puromycin.

### Cloning of ESAG6 3’UTR

RNA was extracted from ∼1 × 10^7^ logarithmic phase cells using the RNeasy plus kit (Qiagen) according to the manufacturer’s instructions. A two-step RT-PCR reaction was performed by first transcribing 0.25 μg of RNA using an Omniscript Reverse Transcriptase (Promega) with a Oligo-dT adapter primer (5’-CGCGTCGACTAGTACTTTTTTTTTTTTTTTT-3’) and then using 5 μl of the resulting cDNA as a template for a PCR amplification using Hot-start RED-Taq (Sigma) with primers specific for BES1 *ESAG6* ORF (6)(5’-GCAGTACATTTGAGTCTT T-3’) and the adapter sequence (5’-CGCGTCGACTAGTAC-3’). The resulting amplicon was ligated into pGEM-T-easy (Promega) prior to DNA sequencing.

### Generation of reporter cell lines

The pRPaΔ^GFP^ X vector, a version of the pRPa^GFP^ X vector (18) with tetracycline operator removed, was a kind gift from Sam Alsford, LSTHM. The firefly Luciferase (fLUC) ORF was PCR amplified from the pCRm-LUC-HYG vector (20) (a kind gift from Phillip Yates, Oregon Health & Science University, USA) using a mutagenic forward primer incorporating a *Hind*III site (underlined) to removes an internal *ApaI* site (mismatch in bold) (5’- ATTAT<u>AAGCTT</u>ATGGAAGATGCCAAAAACATTAAGAA**A**GGCCCAGGG-3’) and a reverse primer incorporating a *Bam*HI site (underlined) (5’-TTCG<u>CGGATCC</u>TCACACGGCGATCTTGC-3’). The PCR product was ligated into pGEM-T-easy (Promega) to allow DNA sequencing prior to subsequent subcloning into pRPaΔ^GFP^ X using the *Hind*III and *Bam*HI sites to replace the GFP- stuffer protein fusion, resulting in pRPaΔ^*fLUC*^ that contains the *fLUC* gene fused to the aldolase 3’UTR.

The 335bp *ESAG6*3’UTR was PCR amplified from *T. brucei*gDNA using a forward primer incorporating *Xba*I (italics) and *Bam*HI (underlined) sites (5’-AATGATCTAGATAG<u>GGATCC</u>GGGAAGGATGCGAC-3’) and reverse primer incorporating an ApaI (underlined) site (5’-AAT <u>AGGGCCC</u>AGTAGAATTAGTCTAGTTT-3’). Digestion with *Xba*I and *Apa*I allowed subcloning into pRPaΔ^GFP^ X, replacing the stuffer protein X and creating pRPaΔ^*GFP*^*-ESAG6-3’UTR* where the *GFP* gene is fused to *ESAG6-3’UTR.* Digestion of the same PCR product with *Bam*HI and ApaI allowed subcloning into pRPaΔ-*fLUC* creating pRPaΔ^*fLUC*^*-ESAG6-3’UTR* where the *fLUC* gene is fused to *ESAG6-3UTR*. Finally, the *ESAG6-3’UTR* of pRPaΔ^*GFP*^*-ESAG6-3’UTR* was replaced with the aldolase 3’UTR of pRPaΔ^GFP^ X using the *Bam*HI and *Apa*I sites creating pRPaΔ-*GFP* that contains the *GFP* gene fused to the aldolase 3’UTR.

The reporter constructs were digested with *Asc*I, ethanol precipitated and resuspended in water prior to transfection. 3 μg DNA was mixed with 1 × 10^7^ 2T1 cells suspended in transfection buffer (21) in a 2 mm cuvette (BioRad) and electroporated in an Amaxa Nucleofector (Lonza Biosciences) using program X-001. Cells were allowed to recover for 6h at 37 °C, 5% CO_2_ in HMI-11T antibiotics before selection with 2.5 μg/mL Hygromycin and 0.5 μg/mL Phleomycin.

### Induction of iron starvation

The iron chelator deferoxamine (Sigma Aldrich) was added to *T. brucei* cell cultures at 25 μM to induce iron starvation. For serum switching starvation experiments, cells were harvested at 800 × g, washed once in HMI-11T without serum, and resuspended in HMI-11T containing either 10% fetal calf serum (Labtech) or 10% donor dog serum (Labtech) with and without the addition of 200 μg/ml holo bovine transferrin (Sigma).

### Western blots

Cells were counted, harvested at 800 × g, the pellet resuspended at 1 × 10^6^ cells/μl in 1 × SDS loading buffer (Melford), and the cells lysed for 5 min at 95 °C. The lysate was separated on a 10% Tris-acetate SDS-PAGE gel, transferred to a PVDF membrane using a Trans blot Turbo semi-dry blotter (Biorad) and blocked overnight in 5% w/w ECL blocking agent (GE Healthcare) in TTBS (0.05% Tween 20 in Tris buffered saline). The expression of GFP was detected with a rabbit anti-GFP primary (1:1000, Roche) and an anti-rabbit-HRP conjugated secondary (1:20,000, GE Healthcare) with expression level quantified with Clarity ECL chemiluminescent detection (Biorad) in a ChemiDoc XPS+ system (Biorad). The detection of tubulin using mouse anti-tubulin KMX-1 primary (1:100; Gift from Keith Gull, Oxford) and anti-mouse-HRP secondary (1:20,000, GE Healthcare) was used to verify loading

### Luciferase assays

The expression of the fLUC reporter gene was quantified by incubating 100 μl of a 5 × 10^5^ cells/ml *T. brucei* culture in white 96 well plates with an equal volume the OneGlo luciferase reagent (Promega) for 5 min, with the resulting luminescence detected using a Fluoroskan ascent FL plate reader (Thermo Scientific) with a 10 sec acquisition time per well.

### Immunofluorescence microscopy

Approximately 1 × 10^6^ cells were harvested by centrifugation at 800 × g, washed once with 1 × PBS and allowed to adhere to glass slides for 5 min. Slides were incubated with 4% PFA for 5 min at room temperature, washed twice in 1 × PBS for 5 minute. A drop of Fluoroshield with DAPI (Sigma) was added to the slide, a coverslip was applied and the slide examined on a Leica DMRXA2 fluorescent microscope.

## Results

### Identification of the ESAG6-3’UTR

In order to investigate whether the 3’UTR of the *Tb*TfR is involved in its dynamic regulation it was necessary to first identify the size of the 3’UTR, which due to its telomeric location is not present in the previous polyadenylation site mappings (22, 23). As expression of ESAG7 may not be essential (6), we focussed our efforts on ESAG6. To identify the size of the *ESAG6 3’UTR* expressed in the active BES1, a two-step RT-PCR was performed with an oligo-dT adapter primer to transcribe cDNA and subsequent amplification of the *ESAG6-3’UTR* with primers specific for BES1 *ESAG6* open reading frame and the adapter sequence. Sequencing of the resulting amplicons identified that polyadenylation occurred at closely spaced sites 317 bp, 330 bp or 335 bp downstream of the *ESAG6* stop codon. The observed heterogeneity and 3’UTR length are consistent with previous observations in *T. brucei* (22, 23). Sequence alignment revealed a high level of sequence conservation between the 13 copies of the *ESAG6-3’UTR* found in the 15 different BES (6), with pairwise comparison of the nucleotide sequences revealing 91.0 – 99.7 % identity (24) (Fig 1 & S1 Fig). Interestingly the 3’UTR of the putative genomic copy of ESAG6 (Tb927.9.15680) was less conserved, with only 38.9 – 42.2 % sequence identity with the telemeric sequences (24) (S2 Fig).

**Fig 1.**
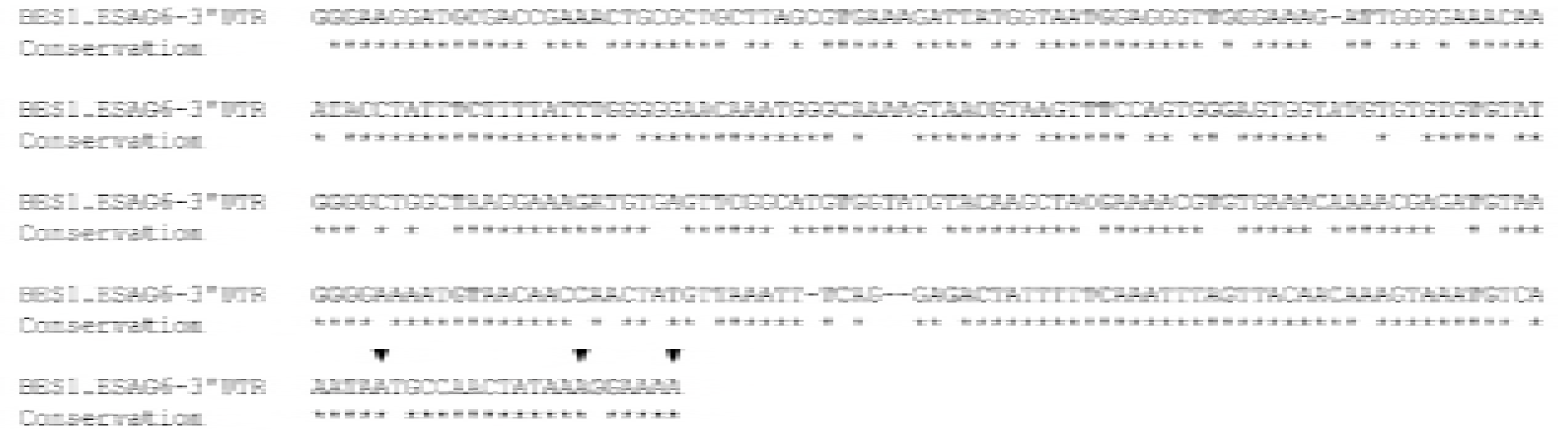
Identification of the *ESAG6-3’UTR*. The observed polyadenylation sites are indicated with triangles above the sequence, and sequence identify with the other 13 BES *ESAG6-3’*UTR sequences indicated by stars below the sequence. Sequences were aligned with T-Coffee (25).

### Fusion of the ESAG6-3’UTR represses reporter protein expression

To investigate the potential involvement of the *ESAG6*-3’UTR in regulation of the *Tb*TfR, constructs were created with the reporter genes *GFP* or firefly Luciferase (*fLUC*) flanked by either the aldolase (*ALD*) 3’UTR or the 335 bp *ESAG6*-3’UTR (Fig 2). The reporter genes were located downstream of a RRNA promoter and integrated into the tagged *RRNA* locus in the 2T1 cell line to ensure high levels of constitutive transcription by RNA pol I whilst avoiding positional effects (18).

**Fig 2.**
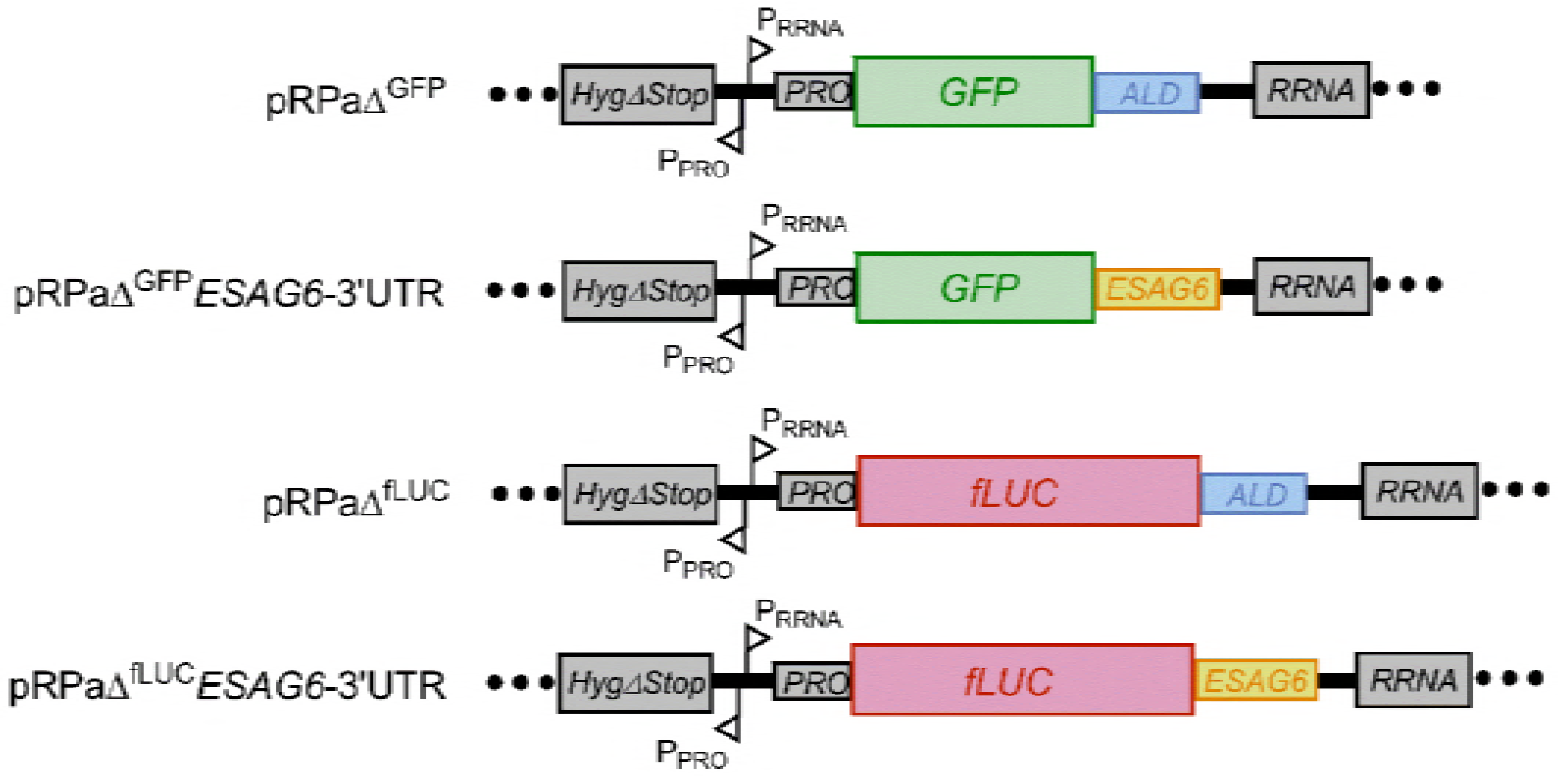
Reporter constructs used in this study. RNA processing regions and promoters: *HygΔStop* - portion of the hygromycin resistance targets the Hyg-tagged RRNA locus; P_RRNA_ - RRNA promoter; P_PRO_-procyclin promoter; *PRO* - procyclin 5’UTR; *ALD* - Aldolase 3’UTR; *ESAG6 - ESAG6-*3’UTR; GFP – green fluorescent protein; fLUC – firefly luciferase.

Observation of the GFP reporter cell lines by immunofluorescence microscopy confirmed cytosolic expression of GFP, with the *ESAG6*-3’UTR reporter cells producing noticeable diminished GFP signal compared to the *ALD*-3’UTR reporter cells (Fig 3A). The steady state expression level of GFP during exponential growth was quantified by western blotting with anti-GFP antibodies for three independent clones, revealing that expression of GFP in the *ESAG6*-3’UTR reporter cells was repressed by ∼80% compared to the *ALD*-3’UTR reporter cells (Fig 3B). Similarly, measurement of luciferase activity in the fLUC expressing cell lines revealed a significant difference between the *ESAG6*-3’UTR and *ALD*-3’UTR for each of three independent clones, with the fusion of the *ESAG6*-3’UTR repressing the expression level by ∼90% (Fig 3C). The level of expression of *Tb*TfR has previously been reported to increase up to 5-fold at high cell density (26), and we observe similar cell density dependent changes in luciferase activity in the *ESAG6-*3’UTR fusion (Fig 3D). Together, these data demonstrate that fusion of the *ESAG6*-3’UTR to a reporter gene is sufficient to repress the expression of the reporter under normal growth conditions.

**Fig 3.**
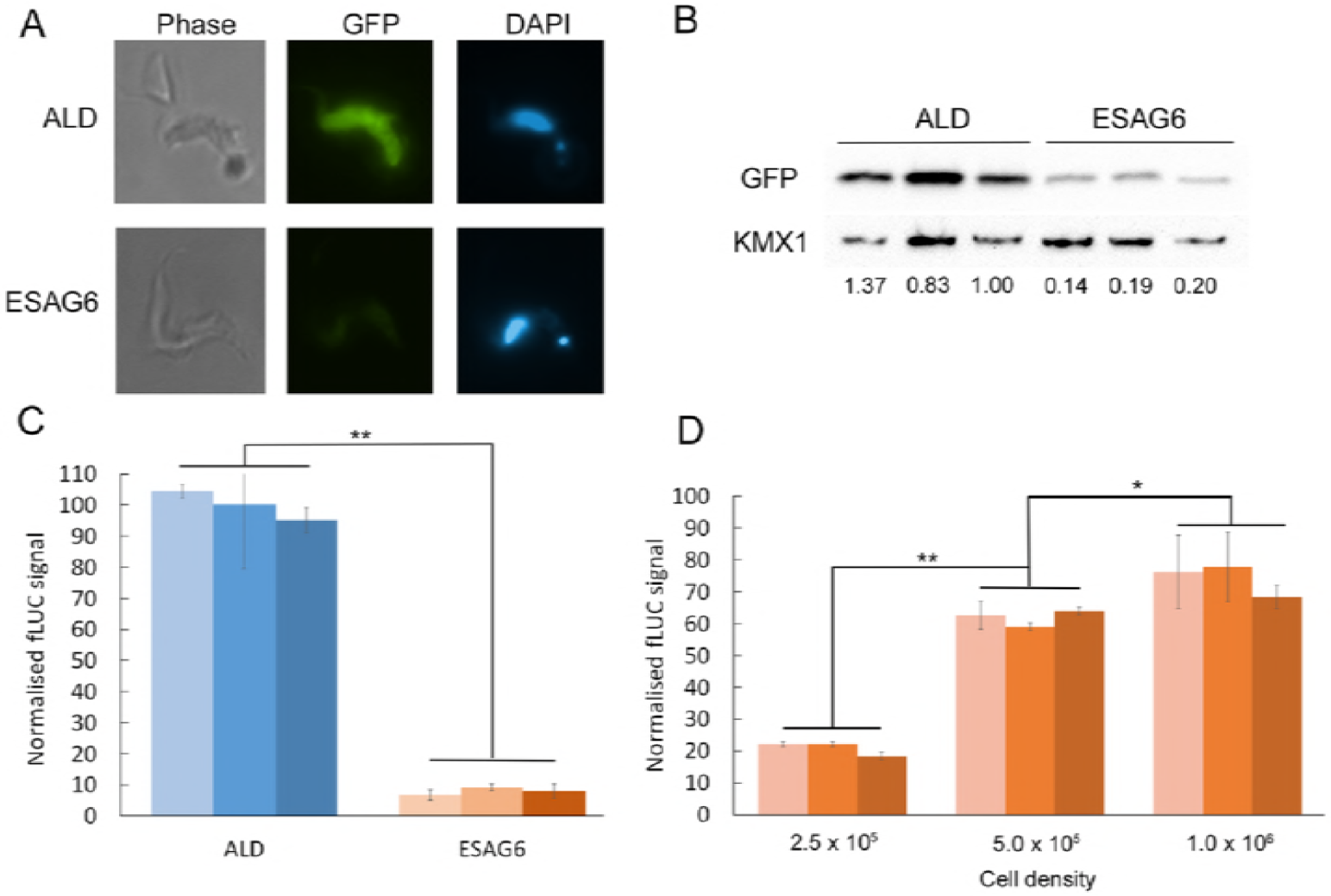
Reporter protein expression under normal culture conditions. A. Immunofluorescence of GFP reporter constructs. B. Quantification of GFP expression by western blotting, using anti-GFP and KMX1 as a loading control. C. Luciferase activity assay, D. Luciferase activity of the *ESAG6*-3’UTR cell line varies with cell density. Data in panel B-D represents three independent clones of each cell line. Luciferase signal is normalised to the aldolase-3’UTR signal; error bars are SEM for triplicate measurements. ALD – aldolase-3’UTR, ESAG6 – ESAG6-3’UTR. * *p* < 0.05, ** *p* < 0.001.

### The ESAG6 3’UTR mediates iron starvation response

The addition of the iron chelator deferoxamine to *T. brucei* cells reduces the availability of Tf-bound iron, and results in a 2.4 – 3.7-fold increase in *Tb*TfR expression after 5 hours (9). Treatment of the fLUC reporter cell lines with deferoxamine for 5 hours had no significant effect on the luciferase activity of the *ALD-*3’UTR reporter cells (Fig 4A). Conversely, deferoxamine treatment increased the luciferase activity of the *ESAG6-*3’UTR reporter cells by ∼10-fold, so that the activity was approaching that of the *ALD-*3’UTR reporter cells (Fig 4A).

**Fig 4.**
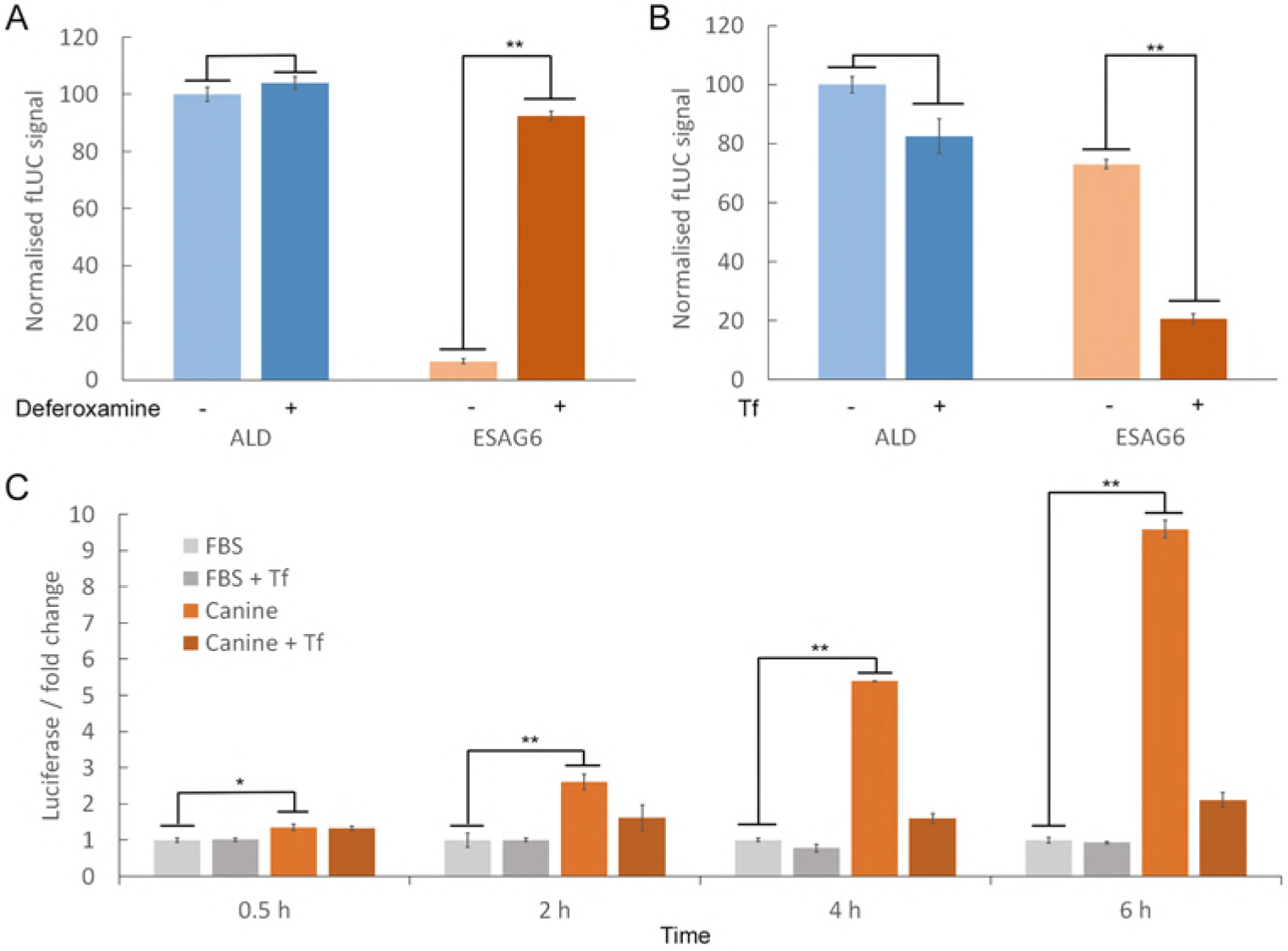
Luciferase expression in the *ESAG6*-3’UTR reporter increases under iron starvation conditions. A. Luciferase activity of *fLUC ESAG6-*3’UTR reporter cells treated with 25 μM deferoxamine for 5 h, normalised to untreated ALD signal. B. Luciferase activity of *fLUC ESAG6-*3’UTR reporter cells switched to HMI11-T + 10% dog serum for 5 h with or without the addition of 200 μg/ml bovine transferrin (Tf), normalised to untreated ALD signal. C. Time course of luciferase activity of *fLUC ESAG6-*3’UTR reporter cells switched to HMI11-T + 10% dog serum for 5 h with or without the addition of 200 μg/ml bovine Tf, normalised to signal in HMI11-T + 10% FCS. ALD - aldolase 3’UTR, ESAG6 - ESAG6 3’UTR. Error bars are SEM for triplicate measurements, * *p* < 0.05, ** *p* < 0.001.

Switching *T. brucei* cells with an active BES1 from growth in media supplemented with 10% bovine serum to growth in media supplemented with 10% canine serum results in rapid 2.5 – 5-fold upregulation of *ESAG6* and *ESAG7* at the mRNA and protein level, as the *Tb*TfR expressed from BES1 has a particularly low affinity for canine Tf (3, 9, 10, 26). The addition of excess bovine Tf to media supplemented with 10% canine serum prevents the upregulation of *Tb*TfR as cells are no longer starved of iron, validating the specificity of the response (3, 4). Switching the fLUC reporter cell lines to canine serum supplemented media for 5 hours increased the luciferase activity of the *ESAG6-*3’UTR reporter cells by ∼3.5-fold compared to growth in canine serum supplemented media with excess bovine Tf (Fig 4B). The addition of excess bovine Tf to bovine serum supplemented media for 5 hours had no significant effect on the luciferase activity of either reporter cell line (S3 Fig).

The *fLUC ESAG6-*3’UTR reporter cell line was used to investigate the temporal profile of the response to serum switching. In the absence of additional bovine Tf, a time-dependent increase in luciferase activity was observed in cells grown in canine serum supplemented media compared to cells grown in bovine serum supplemented media (Fig 4C), with a significant 1.4-fold increase observed as early as 30 minutes after serum switching and increasing to ∼10-fold after 6 hours. Addition of bovine Tf to the canine serum supplemented media reduced the magnitude of the increase to ∼2-fold after 6 hours but did not completely negate the effect, suggesting that the amount of additional bovine Tf used here is not sufficient to completely restore iron intake.

These data demonstrate that fusion of the *ESAG6-*3’UTR to a reporter gene is sufficient to confer dynamic regulation of reporter expression in response to iron starvation conditions, and that upregulation is more rapid than previously thought. The changes in the luciferase activity observed here are broadly consistent with the magnitude of increase in TfR expression previously observed for the iron starvation response, with levels as high as 10-fold reported (26, 27). The slightly larger changes observed with the fLUC reporter compared *Tb*TfR might be due to the superior quantitation and dynamic range of the bioluminescence assay compared to western blotting with polyclonal anti-*Tb*TfR antibodies.

## Discussion

Our data demonstrate that fusion of the *ESAG6-*3’UTR to reporter genes is sufficient to confer a specific response to iron starvation conditions that increases the expression of the reporter with a magnitude and temporal profile consistent with that previously observed for *Tb*TfR upregulation. Since the reporter gene is located in a *RRNA* locus, this provides evidence that the dynamic regulation of *Tb*TfR expression is independent of transcription within the BES site or location at the telomere, and decouples the rapid dynamic regulation of *Tb*TfR expression from subsequent VSG switching events (3, 4, 11). To our knowledge, this is the first time that a *T. brucei* 3’UTR has been demonstrated to be involved in dynamic post-transcriptional regulation of a gene in response to a specific nutritional stimulus, rather than mediated through irreversible developmental regulation in response to lifecycle changes.

We have demonstrated that fusion of the *ESAG6* 3’-UTR causes the expression of the gene product to be repressed under basal conditions, which is consistent with the low level *ESAG6* mRNA present despite being located on the highly transcribed *VSG* polycistronic transcriptional unit. Under iron starvation conditions the repressive effect of the *ESAG6* 3’UTR is reduced, resulting in an increase of expression that is most likely driven by an increase in transcript abundance, as *Tb*TfR increases equally at the mRNA and protein level (9, 10). The involvement of the 3’UTR in iron starvation response suggest that it contains secondary structural elements or sequence motifs that are recognised by RNA binding protein(s) (RBPs), but the high level of sequence conservation in the UTR makes it likely that any such features are more extensive than the 16-mer regulatory sequence identified in VSG mRNAs (28). Further experiments will be required to identify any regulatory features present in the ESAG6 3’-UTR, and whether they have a positive or negative regulatory role.

*Tb*TfR is a multi-gene family that occurs in a promoter proximal position in the 15 BES, and Ansorge *et al*. have previously used RT-PCR analysis to propose that ∼20% of *ESAG6* mRNA is transcribed from the 14 silent BES due promoter-proximal de-repression of silencing (29). Recent analysis of the surface proteome in culture adapted monomorphic cell line by Ghadella *et al*. (30) detected VSGs and ESAGs from multiple BES in addition to the major active BES, which the authors suggest is due to the occurrence of low abundance of a subpopulation of cells that have switched their active BES. Analysis of our own global proteomic data sets (19, 31, 32) supports their conclusion, as although the most abundant ESAGs detected correlate with the BES of the most abundant VSG, several other VSGs from ‘silent’ BES are detected at lower abundance along with their corresponding ESAGs. Therefore we propose that the transcription of *Tb*TfR from supposedly ‘silent’ BES can at least in part be explained by the occurrence of a low abundance of subpopulation of cells that have switched their active BES, and that any contribution of de-repression of silent BES to the rapid dynamic regulation of *Tb*TfR expression is likely to be minimal.

## Conclusions

Taken together, the data presented here demonstrate that fusion of the *ESAG6-*3’UTR to a reporter gene is sufficient to confer a specific response to iron starvation conditions that increases the expression of the reporter with a magnitude and temporal profile consistent with that previously observed for *Tb*TfR upregulation. The effect is independent from transcription within the expression site or location at the telomere, and is therefore decoupled from VSG switching events. The *fLUC-ESAG6*-3’UTR reporter system is experimentally tractable and provides a simple and reproducible signal-and-response system in cultured monomorphic cells with which to elucidate a signalling pathway essential for the clinically-relevant bloodstream form parasite

## Author Contribution

CB carried out the investigation, participated in formal analysis and helped with writing the original draft; WL carried out the investigation and sequence alignments; NF carried the investigation and participated in formal analysis, ACG & HJB carried out the investigation, MDU conceived and the study, conducted formal analysis, and helped with writing the original draft, and it review & editing. All authors gave final approval for publication.

## Acknowledgments

We thank Dr. Sam Alsford (London School of Hygiene and Tropical Medicine, UK) for providing the pRPaΔ^GFP^ X vector, Prof Phillip A. Yates (Oregon Health & Science University, USA) for providing the pCRm-LUC-HYG vector, and Prof. Keith Gull (School of Pathology, Oxford, UK) for providing the KMX-1 antibody.

## Supporting Information

**S1 Figure.**
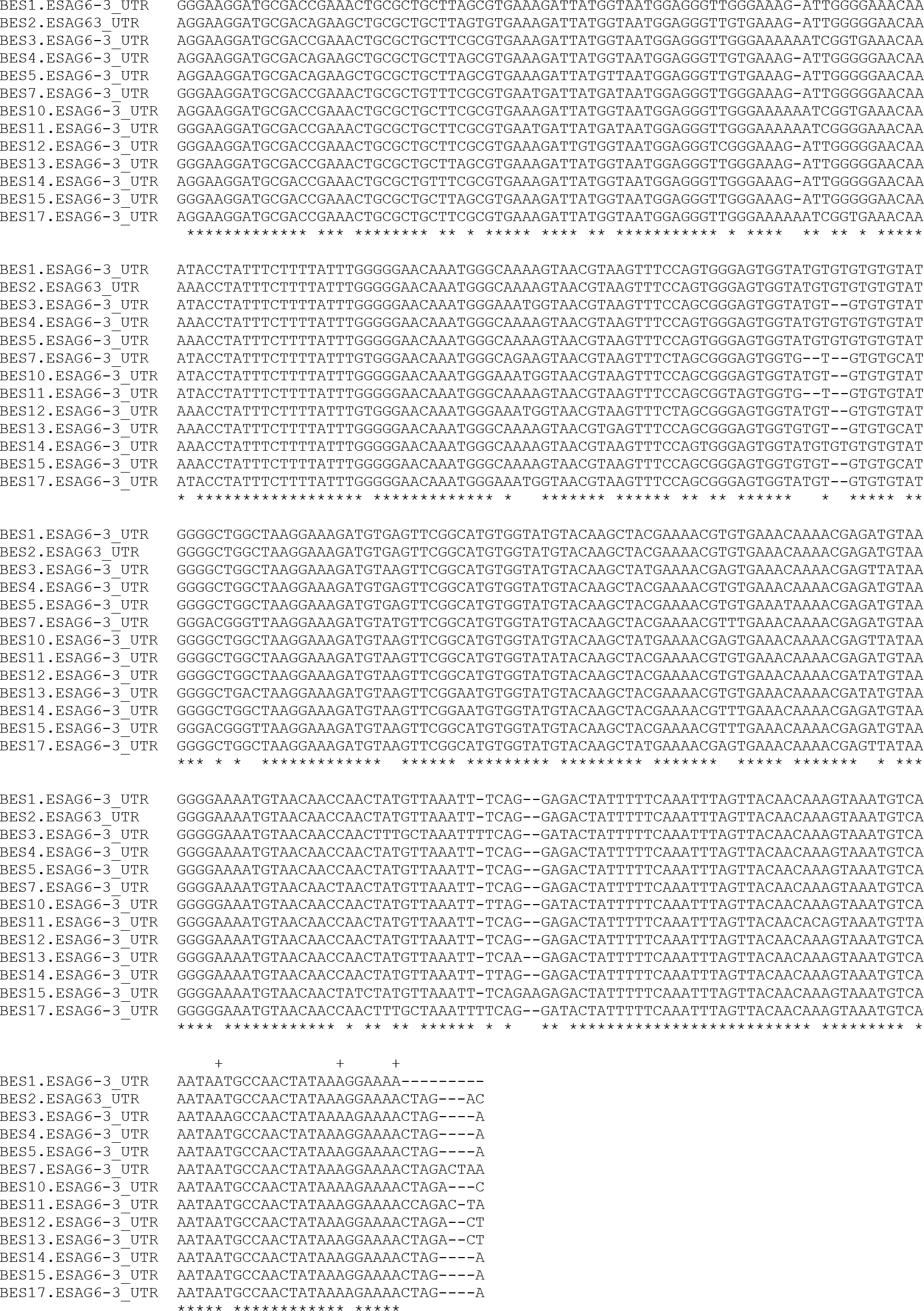
Sequence alignment of the BES ESAG6 3’UTRs. The observed polyadenylation sites are indicated with triangles above the sequence, and sequence identify indicated by stars below the sequence. Sequences were aligned with T-Coffee [1].

**S2 Fig.**
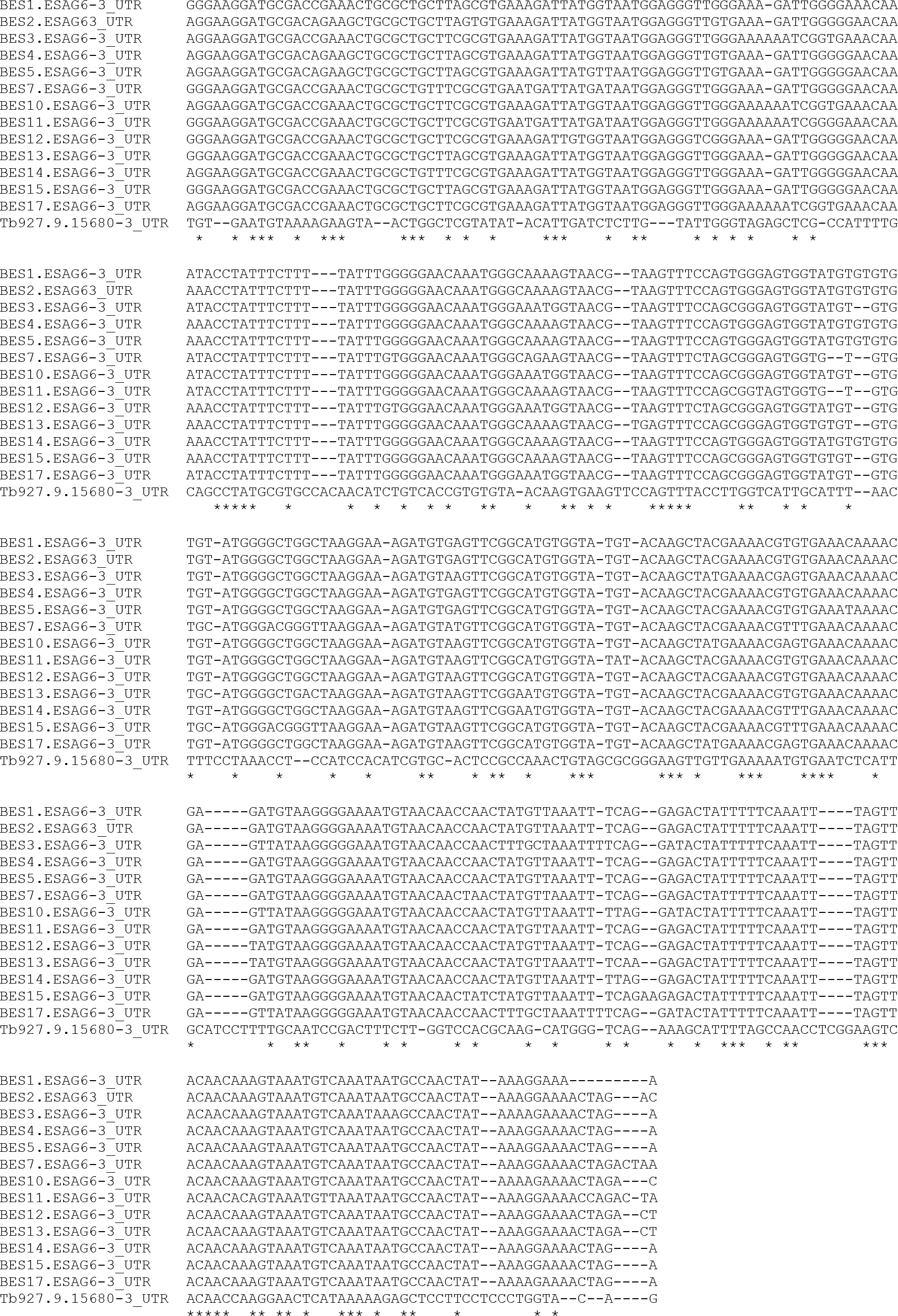
Sequence alignment of the BES and putative genomic ESAG6 3’UTRs. The observed polyadenylation sites are indicated with triangles above the sequence, and sequence identify indicated by stars below the sequence. Sequences were aligned with T-Coffee [1]

**S3 Fig.**
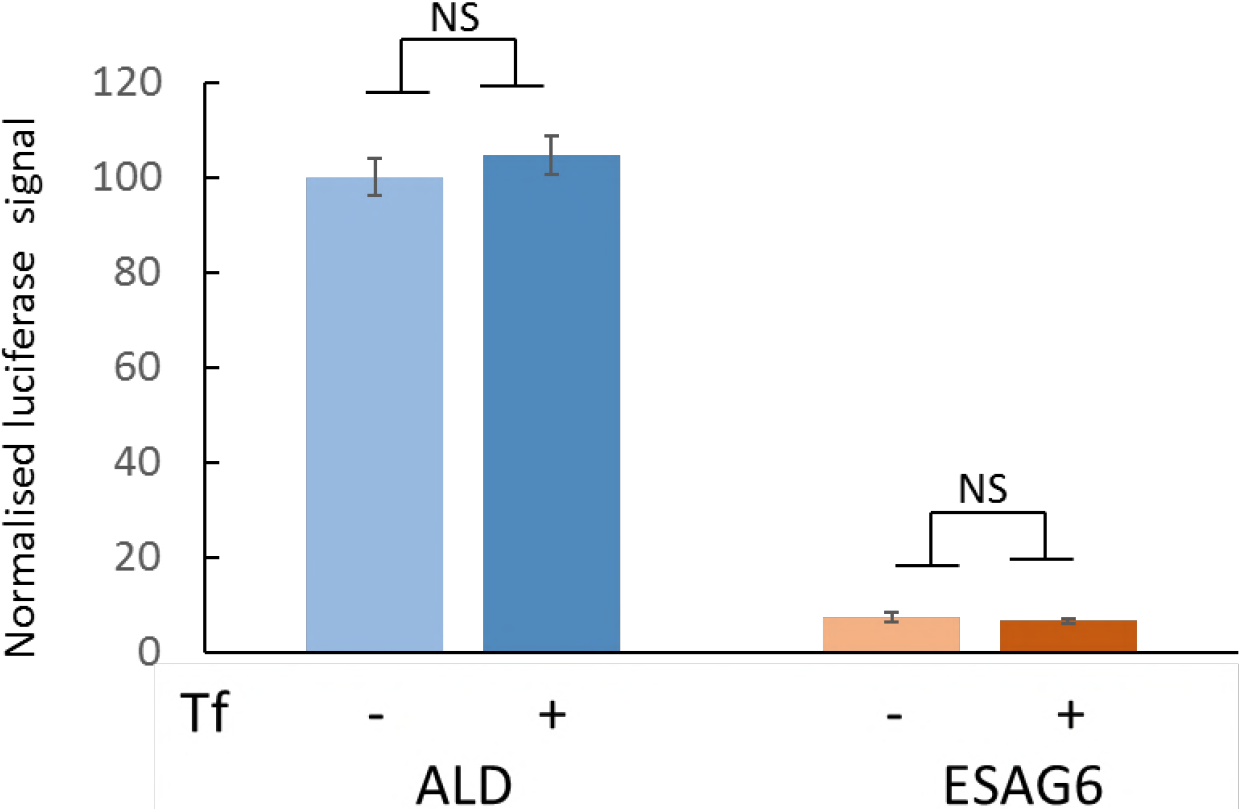
Luciferase reporter expression in normal media is unaffected by addition of bovine transferrin. Luciferase activity of *fLUC ESAG6-*3’UTR reporter cells grown in HMI11-T + 10% FBS with or without the addition of 200 mg/ml bovine transferrin, normalised to ALD signal. ALD – aldolase 3’UTR, ESAG6 – ESAG6 3’UTR. Error bars are SEM for triplicate measurements, NS = *p* > 0.05.

